# Extracting Reproducible Time-Resolved Resting State Networks using Dynamic Mode Decomposition

**DOI:** 10.1101/343061

**Authors:** James M. Kunert-Graf, Kristian M. Eschenburg, David J. Galas, J. Nathan Kutz, Swati D. Rane, Bingni W. Brunton

## Abstract

Resting state networks (RSNs) extracted from functional magnetic resonance imaging (fMRI) scans are believed to reflect the intrinsic organization and network structure of brain regions. Most traditional methods for computing RSNs typically assume these functional networks are static throughout the duration of a scan lasting 5–15 minutes. However, they are known to vary on timescales ranging from seconds to years; in addition, the dynamic properties of RSNs are affected in a wide variety of neurological disorders. Recently, there has been a proliferation of methods for characterizing RSN dynamics, yet it remains a challenge to extract reproducible time-resolved networks. In this paper, we develop a novel method based on dynamic mode decomposition (DMD) to extract networks from short windows of noisy, high-dimensional fMRI data, allowing RSNs from single scans to be resolved robustly at a temporal resolution of seconds. We demonstrate this method on data from 120 individuals from the Human Connectome Project and show that unsupervised clustering of DMD modes discovers RSNs at both the group (gDMD) and the single subject (sDMD) levels. The gDMD modes closely resemble canonical RSNs. Compared to established methods, sDMD modes capture individualized RSN structure that both better resembles the population RSN and better captures subject-level variation. We further leverage this time-resolved sDMD analysis to infer occupancy and transitions among RSNs with high reproducibility. This automated DMD-based method is a powerful tool to characterize spatial and temporal structures of RSNs in individual subjects.

## 1. Introduction

Resting state networks (RSNs) comprise distinct regions of the brain that exhibit synchronous low-frequency (<0.1 Hz) temporal fluctuations in the absence of explicit tasks [1, 2, 3]. They are most commonly detected using blood-oxygen level-dependent (BOLD) resting state functional magnetic resonance imaging (rs-fMRI), where a series of T2 or T2* weighted images of the brain are acquired repeatedly over the duration of the scan (5–15 minutes) [4, 5]. The data-driven extraction of RSNs from these noisy and high-dimensional datasets is a difficult analytic task, made possible through the use of well-suited methods. Independent Component Analysis (ICA) [66] has become a standard approach in this field, due to its power at detecting and disentangling overlapping, but statistically distinct, signals from noisy, high-dimensional data. The traditional application of ICA and other RSN analyses have focused on static analysis of the images, implicitly assuming that the networks are unaltered throughout the duration of each resting-state scan [5]. The assumption that brain states are static for many minutes has the effect of averaging over the spatial and temporal variability of networks [6].

Although the spatial structure of these RSN patterns is similar throughout the population, their exact structure and dynamics in time vary considerably between individuals [7, 8, 9, 10]. Further, even within individuals, the modes reconfigure dynamically on timescales ranging from seconds to years [11, 12, 13, 14, 15, 16]. Differences in an individual’s RSN dynamics are important in part because they may serve as useful biomarkers for a variety of neurological dysfunctions, including schizophrenia [17, 18, 19, 20, 21], bipolar disorder [20, 21], autism [22, 23], depression [24], post-traumatic stress disorder [25], attention deficit/hyperactivity disorder [26], and mild cognitive impairment [27]. For example, multiple studies have shown that in schizophrenia, regions of default mode, auditory, motor, and visual resting state networks show differences in their correlations, when compared to a control population [28, 29, 30]. Patients with schizophrenia also show abnormalities in dynamic dwell times in large-scale RSNs [31].

Technological innovations in brain imaging such as SENSitivity Encoding (SENSE) and simultaneous multi-slice (SMS) acquisitions now allow for improved temporal resolution, enabling investigation of the true temporal dynamics of brain function [32, 33, 34]. One emerging challenge with the advent of the fast acquisition of brain dynamics is the appropriate choice of methods to analyze and interpret data. It follows that much intense work has gone beyond static analyses, calculating RSN structure and activation as they vary in time to investigate a variety of measures of RSN dynamics. Many of these time-resolved RSN analyses use a windowed version of a traditional method, such as correlation [35, 36, 15, 37, 38, 6, 39] or ICA [40, 41]. Windows are then shifted in time and the measures recalculated, leading to methods for characterizing dynamic connectivity [42, 43, 35, 44]. These most common RSN extraction methods make no explicit assumptions about the intrinsic temporal structure of the fMRI data [45, 46]. We note that there are alternatives to the windowed methods such as the wavelet coherence transform approach, which performs a time-frequency decomposition of the resting state signals from a pair of voxels or regions of interest [47]. Another non-windowed approach is to fit a hidden Markov model, which uses Bayesian inference to infer states and their dynamics with the assumption that the system is in precisely one state at any given time [48].

In this work, we present a new framework based on dynamic mode decomposition (DMD) for analyzing resting state BOLD fMRI data. DMD is a spatiotemporal modal decomposition technique [49, 50] ideally suited to extract coherent modes from rs-fMRI data. Similar to ICA, DMD decomposes a signal into constituent coupled spatiotemporal modes [51, 52, 53]. Unlike ICA, DMD constrains the modes to be temporally coherent; specifically, each mode oscillates at a fixed frequency while exponentially growing or decaying. This temporal coherence constraint produces more robust estimates of spatial modes by leveraging the assumption that RSNs have coherent dynamics within short windows of time, while automatically allowing for any phase differences between regions of a network. Thus, in addition to a spatial map of activation, DMD also estimates the temporal frequencies associated with each spatial mode.

Here we develop and validate a novel method to extract reproducible, time-resolved RSNs based on DMD. To demonstrate our method, we apply it to rs-fMRI data from 120 individuals in the Human Connectome Project (HCP) dataset [54, 55, 56]. DMD is computed for rs-fMRI data in windows, and the spatial components of the DMD, known as DMD modes, represent coherent spatial correlations across brain areas (Fig. 1A). We first consider group-DMD (gDMD), where hierarchical clustering discovers stereotypical modes present within the full population (Fig. 1B). These modes are strongly clustered, with clusters and sub-clusters corresponding directly to canonical RSNs. We next applied our DMD analysis independently to single scans (sDMD), as illustrated in Fig. 1C. Our technique extracts and characterizes individualized RSN modes and their dynamics at a resolution of ~3s. To assess our time-resolved results, we compute the subject-level default mode networks (DMNs) through DMD and find that the individualized variations we observe agree with those derived through group-ICA (gICA) and dual regression. Importantly, this agreement is in spite of the fact that our DMD approach uses only data from a single subject, whereas the gICA approach regresses against a reference RSN computed from the entire population. We show that the sDMD modes resemble the gICA modes more closely and more robustly than subject-level DMNs computed via single-scan spatial ICA. Our subject-level analysis lends well to characterizing the RSN dynamics, including how frequently each RSN is active and the probabilities of transitions between different RSNs. We show that the sDMD derived RSN occupancies are reproducible within each individual across independent scans, and these dynamical properties may be used to build subject-level network representations of RSN dynamics. These results lay the groundwork for the application of the DMD family of methods to fMRI data; DMD is a modular approach that extends easily to multi-resolution analyses and control theoretic frameworks, a key advantage over ICA-based methods that have no directly analogous extensions. We suggest that DMD is an improved method for robust, reproducible extraction of functional connectivity modes from short windows of rs-fMRI data, and these time-resolved modes are ideally suited to characterize RSN dynamics on the single-subject level.

**Figure 1:**
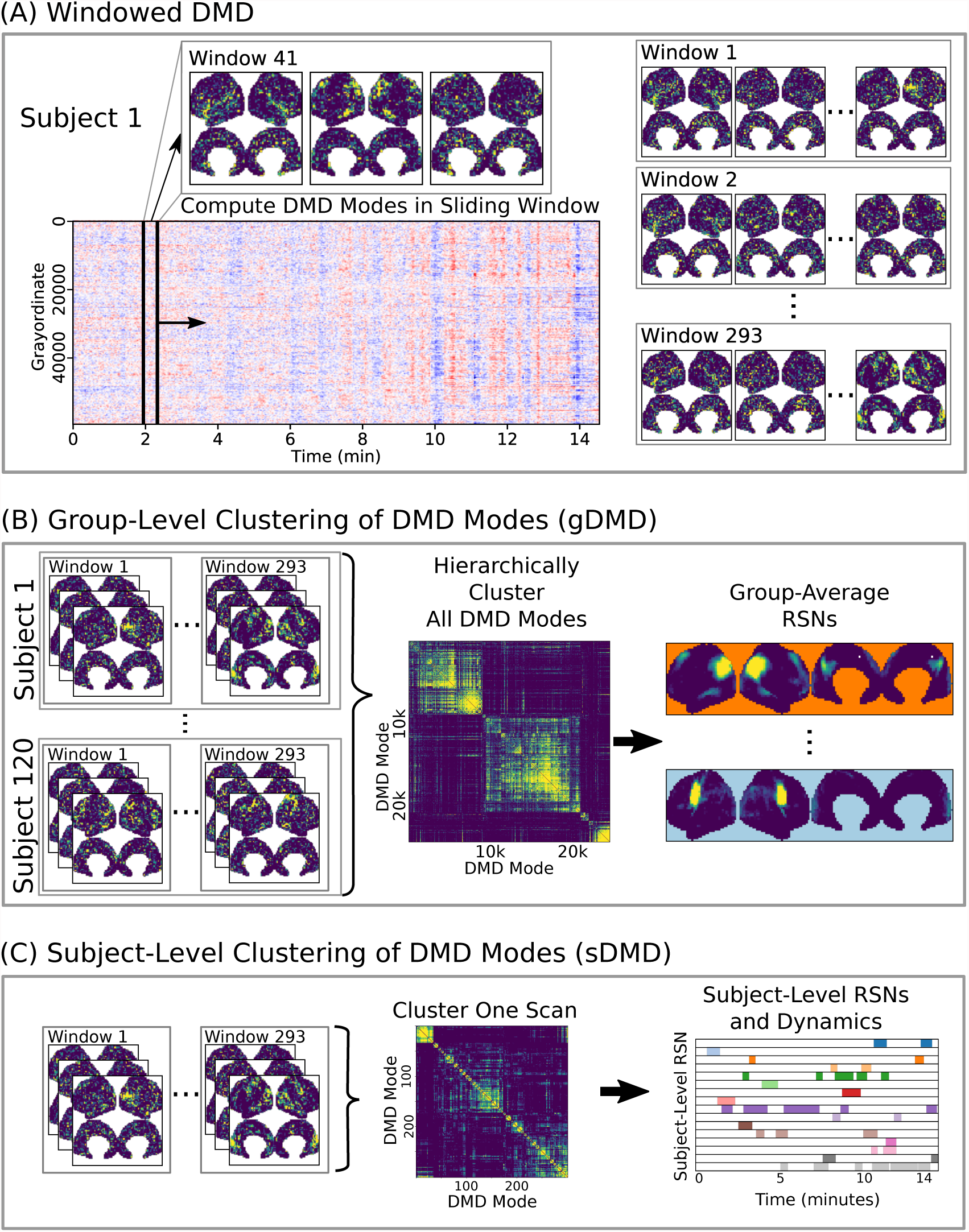
A schematic of our method using DMD to analyze resting state BOLD fMRI data. **(A)** Spatiotemporal DMD modes are extracted from short sliding windows of rs-fMRI data from 120 subjects from the Human Connectome Project. We use 23-sec windows and slide in 3-sec steps over each 15-min scan. **(B)** Group-DMD (gDMD) clusters all modes from the full population to extract group-averaged RSNs. **(C)** Subject-level DMD (sDMD) clusters modes from a single scan, which yields both individualized RSNs and their corresponding dynamics.

## 2. Results

Here we present results using our method combining dynamic mode decomposition (DMD) and unsupervised clustering to extract resting state networks (RSNs) from Human Connectome Project (HCP) data. We show results at both the group level (gDMD) and the subject level (sDMD), as summarized in Fig. 1. The sDMD results extract reproducible, time-resolved RSNs and their transition dynamics for individual scans. Details of our approach are described in the Methods. Code to assist in downloading the correct data, run all analyses described in this paper, and generate the corresponding figures are openly available at https://github.com/kunert/DMD RSN.

### 2.1. Extracted group-level modes resemble Resting State Networks (RSNs)

The BOLD data we analyze is a collection of 15-min resting-state scans from 120 unrelated individuals from the HCP. Each scan has 1200 frames acquired at a temporal resolution of ~0.72 seconds. The data had been pre-processed with the HCP minimal pre-processing pipeline [56] and is freely available to download at https://www.humanconnectome.org/. As part of this pre-processing, the data are mapped to the “grayordinate” space, where each point maps onto the cortical surface.

To resolve modes in time, we apply DMD to short windows of data within each individual scan. In the results shown here (Fig. 2), we use ~23-sec windows sliding in ~3-sec increments. This windowing scheme corresponds to windows of 32 frames sliding by 4 frames, producing a total over 293 overlapping windows per each 1200-frame scan. DMD decomposes the data in each window into a set of spatiotemporal modes, and we truncate to the first 8 temporally coherent modes; this truncation produces a low-rank representation of the dominant spatiotemporal patterns in the data. Our results presented here are relatively robust to the number of modes calculated within each window, as described further in Appendix C.

**Figure 2:**
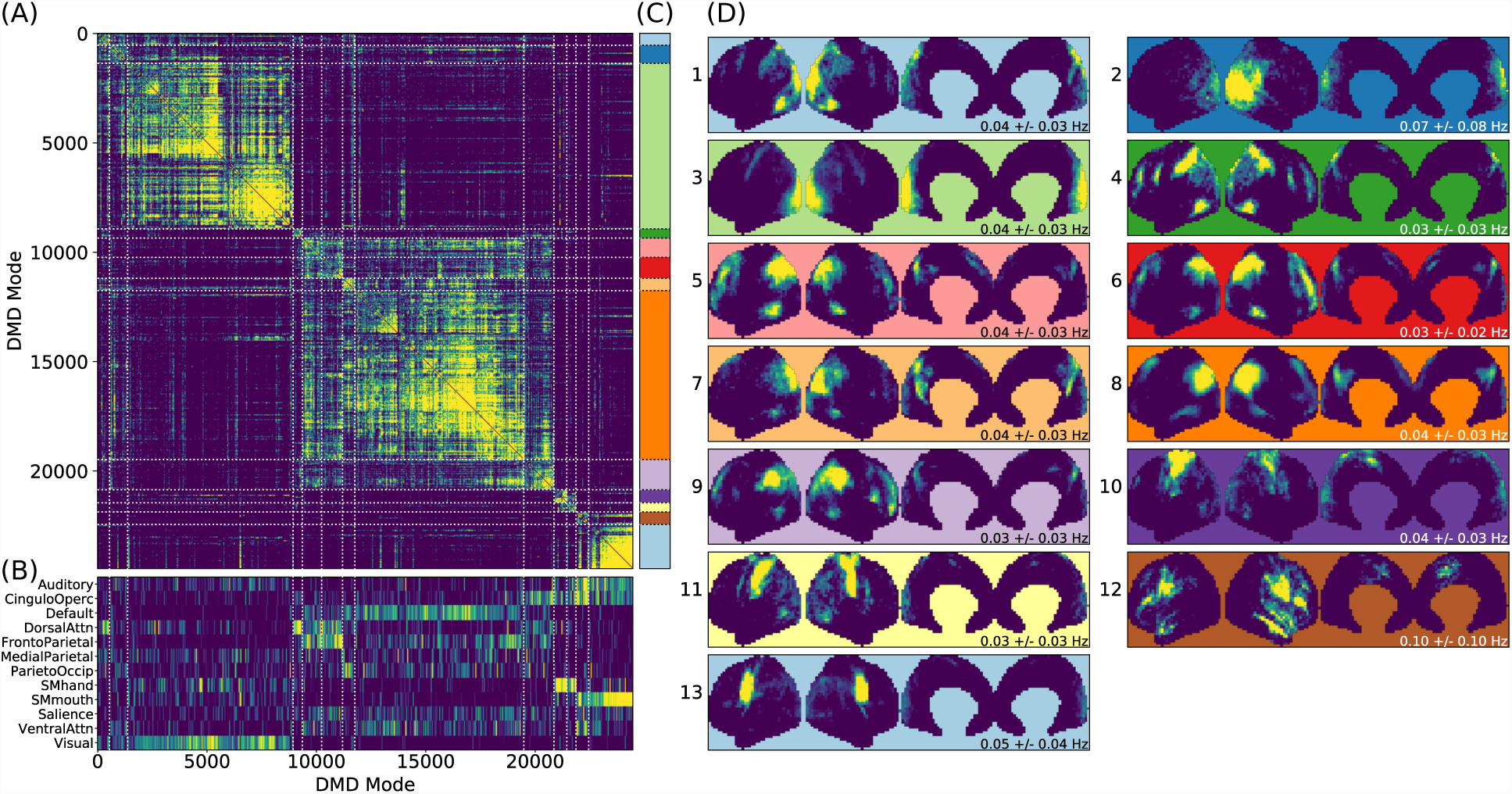
Results from gDMD using modes extracted from scans of 120 individuals, revealing clusters closely resembling known RSNs. **(A)** The hierarchically clustered correlation matrix between gDMD modes. **(B)** Overlap of each mode with RSNs from [57]. **(C)** Clusters automatically extracted from the hierarchical clustering results. A different threshold can be chosen to derive finer or coarser sub-clusters. **(D)** Averaged modes within each of the 13 clusters, with the corresponding DMD temporal frequencies. Several of the automatically identified DMD mode clusters visually correspond to canonical RSNs.

To show that the DMD modes thus computed resemble known RSNs, we use an unsupervised clustering approach to automatically identify and label clusters of modes, then compare them to canonical RSNs. Specifically, we perform agglomerative clustering to hierarchically cluster our set of DMD modes based on the average spatial correlation between clusters [59, 60] (as implemented in the `cluster.hierarchy` package of SciPy 1.0.0 [58]). In gDMD, we consider as a group all the DMD modes from all windows of 120 individuals in the dataset. The hierarchically clustered correlation matrix is shown in Fig. 2A, where the modes have been reordered so that this matrix is strongly block diagonal. In other words, groups of modes cluster naturally. Fig. 2B shows the correlation of each DMD mode to each of a set of reference RSNs [57]. Importantly, the automatically-grouped clusters appear to correspond to distinct RSNs. For instance, the strongly clustered block in the lower-right of the correlation matrix shows very strong correlations to the mouth sensorimotor RSN.

The clustering establishes boundaries that separate modes into distinct clusters, shown in different colors in Fig. 2C. This procedure uses a threshold that determines the resultant size and granularity of the clusters. Here we choose a threshold on the cophenetic distance (i.e. the “distance” threshold option of the `scipy.cluster.hierarchy` function `fclust` [58]) of 0.955, a relatively coarse grouping of the modes into 13 clusters whose averages are shown in Fig. 2D. These cluster averages, extracted at the population level, appear to correctly resemble known resting state networks. For instance, cluster 3 (light green) resembles the visual network, cluster 8 (dark orange) resembles the default network, and cluster 10 (dark purple) resembles the sensorimotor hand network.

Every DMD spatial mode has a corresponding temporal eigenvalue, which gives information about the oscillation frequency of that mode in time. Although these eigenvalues have not been used in the clustering process, they describe the characteristic oscillation frequency of each cluster. The mean and standard deviation for each cluster’s oscillation frequency is shown in Fig. 2D. As a validation that RSNs have low frequencies of oscillation, these frequencies are generally below or around 0.10 Hz. Further analysis of the frequency content of extracted gDMD clusters is discussed in Appendix B.

### 2.2. Subject-level modes and dynamics

One key advantage of our approach using DMD followed by unsupervised clustering is that it can be performed equally well on any subset of modes. We are particularly interested in performing the analysis on single scans, which we refer to as sDMD. Fig. 3 shows the results from hierarchically clustering modes from a single 15-minute scan. Similarly to the group results, this similarity matrix is strongly block diagonal, and the blocks correspond to different canonical RSNs (Fig. 3A, bottom). Note that we use a slightly lower threshold on the cophenetic distance than in Fig. 2 (0.95 instead of 0.955), yielding a larger number of finer-grained subclusters.

**Figure 3:**
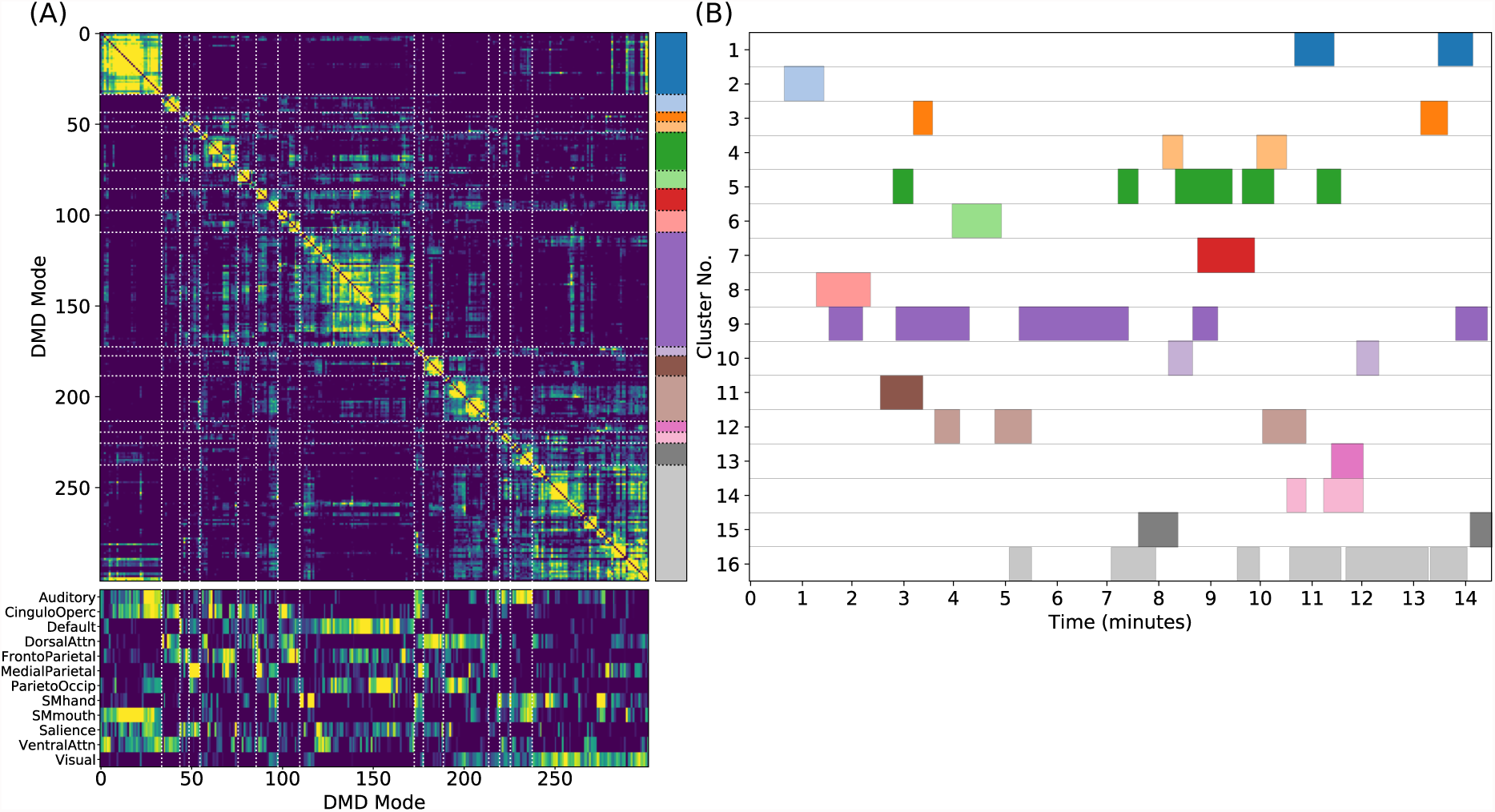
Results from sDMD using modes extracted from a single scan. **(A)** The hierarchically clustered correlation matrix between the sDMD modes, and their overlap with the RSNs of [57]. **(B)** Temporal activation of each cluster within the scan. Each row corresponds to a distinct cluster, and columns are colored in if the corresponding cluster is active within that time window.

Importantly, in this single-scan analysis, the clusters are easily interpreted temporally: each cluster shows temporal dynamics defined by the time windows in which its constituent modes are found, as shown in Fig. 3B. Largely, clusters are active over periods spanning many consecutive windows and extended periods of time. Notably, many different clusters are observed to co-occur in time. This overlap of modes in the same window poses no problems for the sDMD approach, but it violates the assumptions made by other time-resolved methods (such as hidden Markov models) that require the system is in a single state at any particular time.

### 2.3. Subject-level modes capture spatial heterogeneity

Clusters derived by sDMD are able to reliably capture individual variability of RSNs. To quantify this feature, we compare one particular resting state network, the default mode network (DMN), as extracted by three different methods: sDMD, ICA, and group-ICA with dual regression (gICA). The gICA networks were computed by the HCP and are available on ConnectomeDB as part of the *High-level rfMRI Connectivity Analyses* data release. In short, this approach generates high-quality group ICA modes using the dataset of 1200 individuals then uses dual regression to adapt each group mode to the heterogenous structure seen within a particular scan.

Fig. 4 compares the DMNs extracted by ICA, sDMD, and gICA. Specifically, we use spatial-ICA on the entire window of scan data, as opposed to the sliding-window approach of sDMD. ICA and sDMD were both run multiple times with varying hyper-parameters, and for each method we choose the output that most strongly correlates with the canonical DMN. Specifically, ICA was performed multiple times as the fitting process is inherently stochastic, and will sometimes fail to extract a quality mode; thus we performed ICA on each scan ten times, each time randomly varying the number of modes in the range 8-16, and kept the mode which most strongly resembled the gICA DMN. sDMD, on the other hand, is inherently deterministic and was only performed once to produce a single set of modes; clustering of these modes was performed over a range of clustering parameters (the pre-clustering mode mask threshold, and the cophenetic distance threshold for forming clusters) and we kept the result most closely resembling the gICA mode. Importantly, the ICA and sDMD analyses were computed on data from single scans, whereas gICA produces personalized DMNs for each scan based on a population-derived DMN using dual regression. As shown in the examples from 3 different subjects in Fig. 4A, qualitatively speaking, ICA and sDMD are both able to extract DMNs that resemble the personalized gICA DMNs. In particular, note that some of the spatial heterogeneity of DMNs for individual subjects is recapitulated in the ICA and sDMD networks.

**Figure 4:**
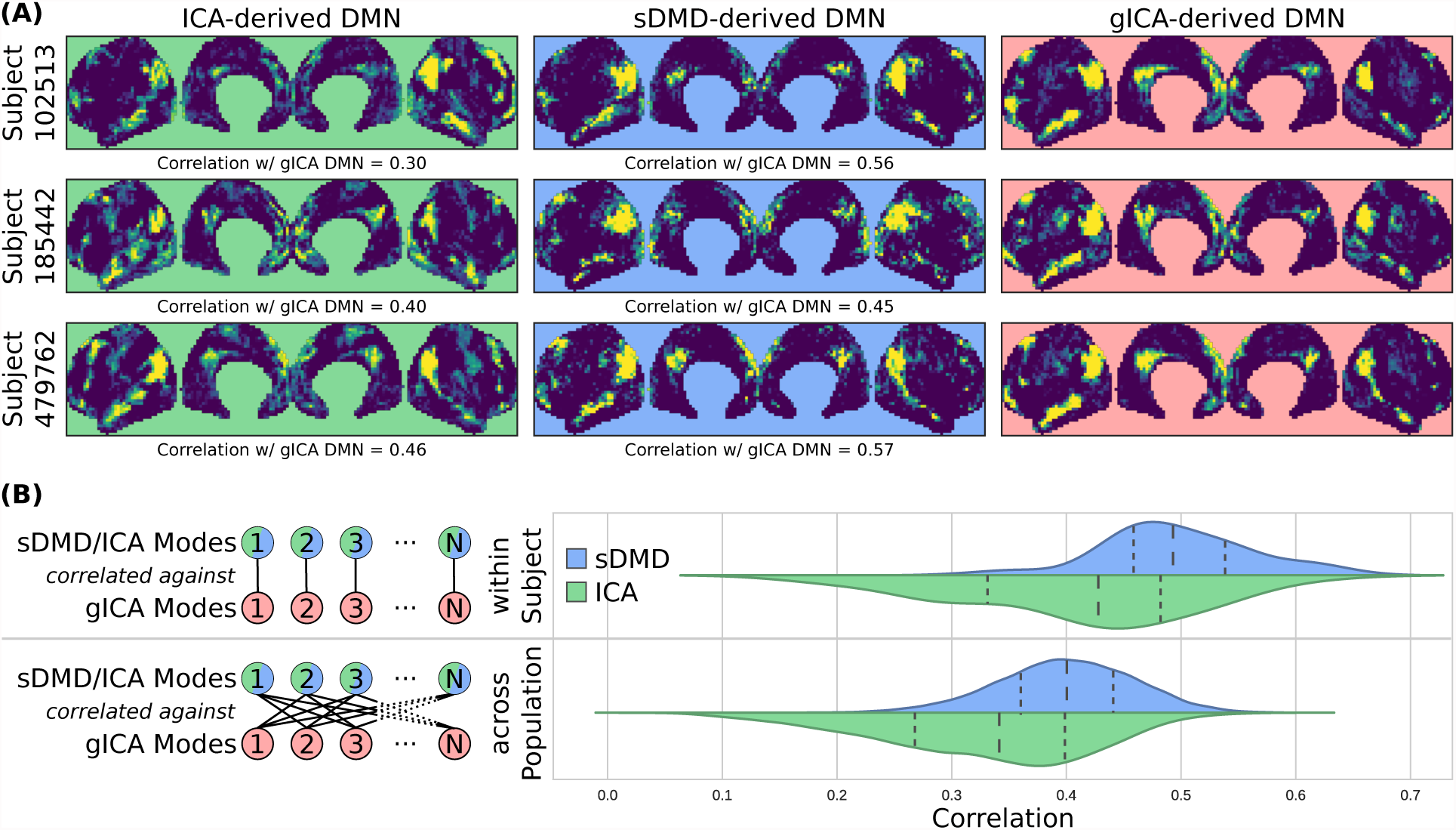
sDMD is able to capture the individual heterogeneity in spatial DMN structure. **(A)** Each column shows examples of subject-level DMNs as calculated by: (i) traditional ICA, (ii) sDMD, and (iii) group-ICA with dual regression (gICA). gICA yields the cleanest modes but requires a high-quality group-level mask, whereas ICA and sDMD use only data from a single scan. **(B)** The spatial correlation of ICA/sDMD modes with gICA modes. We compare how strongly each ICA/sDMD mode correlates with the gICA mode from the same individual, as well as how strongly it correlates with the full population’s gICA modes. Both methods find modes which correlate more strongly with DMNs from the same individual, indicating that they all capture similar subject-level variations in DMN structure. sDMD outperforms ICA in this regard, however, and correlates more strongly with the gICA modes. At the same time, sDMD provides more unambiguous temporal information than either approach.

We show that sDMD reliably outperforms ICA in quantitative comparisons with gICA on a subject-by-subject basis. Fig. 4B shows that that DMNs from both ICA and sDMD correlate significantly more strongly with gICA results from the same individual (bottom panel) than with gICA results from the rest of the population (top panel). Specifically, the distribution of correlations for ICA has a lower mean and longer tail of low correlation values than the distribution of correlations for sDMD. This result indicates that sDMD extracts similar individualized DMN structures as gICA using data from only a single subject. Further, the sDMD analysis provides unambiguous time dynamics of the subject-level modes. The gICA or ICA modes can be correlated against the scan data to yield a measure of activation in time, but this is a continuous measure which requires thresholding and will be significantly nonzero in the case of any overlapping signal or noise. Our clustering approach, on the other hand, unambiguously identifies a cluster as active/inactive within a particular window of time.

### 2.4. Dynamic properties of modes are reproducible

Our proposed gDMD and sDMD analyses produce reproducible dynamics in addition to reliable spatial structures. This reproducibility criterion is critical for interpreting dynamic properties as meaningful reflections of the underlying functional networks. Useful dynamic properties to compute include how often each mode is active, and how often pairs of modes are active together (either simultaneously, or with one mode transitioning into another). To assess reproducibility, we take advantage of the HCP resting state dataset, where each of the 120 subjects underwent four (4) different 15-min resting state scans. Specifically, we repeat the same analysis for each of these four sets of scans separately and quantify to what extent our DMD analysis extracts similar modes and dynamic properties. Here we examine dynamic networks extracted by gDMD, looking at two simple measure of dynamics: the occupancy of and the transfer matrices between networks. Next, we quantify the reproducibility of these dynamics at a single-subject level by using sDMD on each of the 4 scans for each subject.

Modes were paired between the four independent scans based upon their spatial similarity; this matching procedure, as well as visualizations of all extracted modes, is detailed in Appendix A. Fig. 5A shows the six most highly correlated modes. Fig. 5B shows the occupancy matrix *O_ij_* of these 6 modes, where the value in the *i^th^* row and *j^th^* column corresponds to how often modes *i* and *j* are simultaneously active. The diagonal *O_ii_* shows how often each mode *i* is active, and the matrix is symmetric. Next, Fig. 5C shows a transfer matrix *T_ij_,* which contains the probability that, if mode *i* is active, then mode *j* will be active 30 seconds later. This matrix not symmetric, and it encodes rich dynamical information that can be interpreted as a dynamic network model, as shown schematically in Fig. 5D.

**Figure 5:**
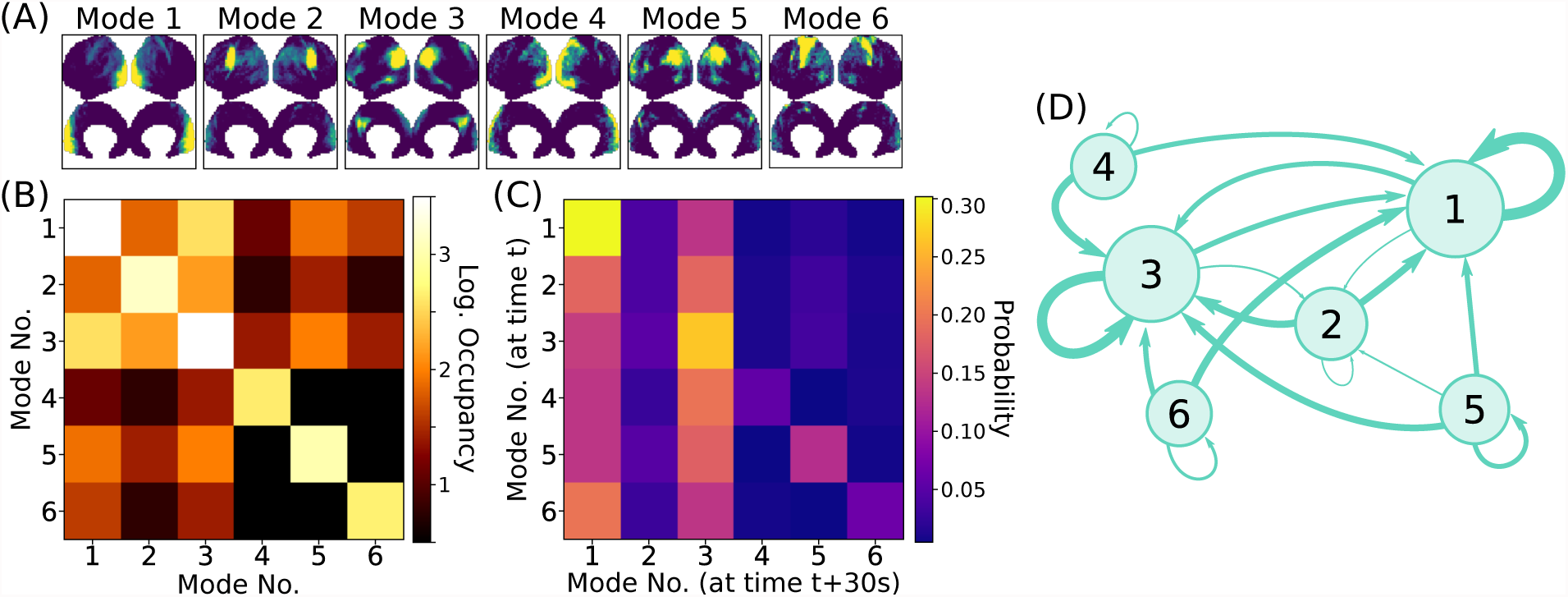
Dynamic properties of networks as extracted by gDMD revealed by the occupancy and transitions of modes. **(A)** gDMD was performed on four independent sets of scans of the same population of 120 individuals. The six modes plotted here are those extracted most reliably from the four datasets. **(B)** The occupancy matrix *O_ij_*: each entry shows how often mode *i* and *j* are active simultaneously (the diagonal simply shows how often mode *i* is active). **(C)** The transfer matrix *T_ij_:* this shows the probability that if mode *i* is active, mode *j* will be active 30 seconds later. **(D)** The transfer matrix can be interpreted as a network which characterizes RSN dynamics.

We next use these same 6 modes from the gDMD and repeat the analysis for individual single scans, analyzing the extent to which the dynamic properties are unique to each individual. The occupancy of each mode was broken up into how often it is present in individual scans, and Fig. 6 shows the results of the subject-level occupancy correlation between two different sets of scans of the same individuals. Correlation coefficients and p-values are calculated for all possible pairs of the 4 different scans, and Fig. 6 reports the median correlation across all scan comparisons and the corresponding p-value. For five of the six modes, occupancy results from different scans of the same individual are positively correlated, with the strongest and most significant correlations observed for the most active modes. Note that the high p-value of Mode 4 does not imply that it is not meaningful, but rather suggest that its activity may not correlate within an individual across different scans (as would be the case, for example, if mode activity was uniformly probable across the population). Thus, our approach extracts reproducible, individualized dynamic properties of resting state networks.

**Figure 6:**
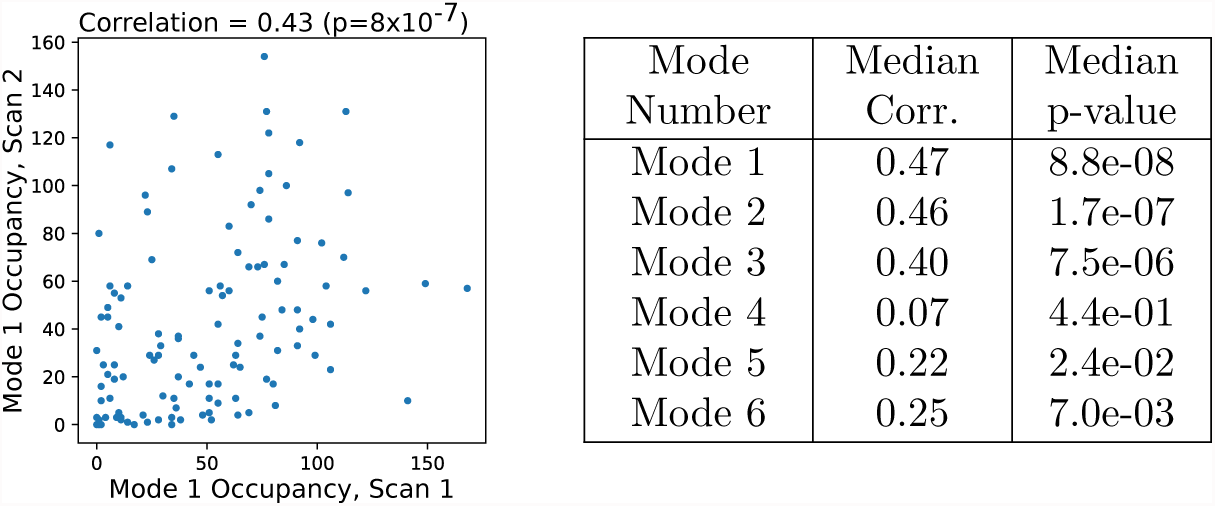
Occupancy of modes computed by sDMD is correlated across different scans of the same subjects. The 6 modes are the same as in those in Fig. 5. Spearman’s rank correlation is computed of mode occupancies between all pairs of the 4 scans. The plot at left shows the occupancy of Mode 1 (*O*_11_) for each subject in two different scans (REST1_LR and REST2_LR). There is a significant correlation; subjects with high activity in one scan are more likely to have high activity in another. In the table at right, we have correlated all possible pairs of the four scans and report the median correlation (and corresponding p-value). The correlations have *p* ≤ 0.05 for all but mode 4.

## 3. Discussion

In this work we present a novel framework based on the dynamic mode decomposition (DMD) for extracting time-resolved resting state networks at both the group (gDMD) and single-subject (sDMD) levels. DMD decomposes high-dimensional time-series data into a sum of dominant coupled spatiotemporal modes. One way to think about DMD is that it is similar to a combination of applying principal component analysis (PCA) in space and the Fourier transform in time, as it extracts both spatial maps and their coupled frequency content simultaneously. We demonstrate our approach using data from 120 subjects from the Human Connectome Project (HCP). DMD to extract coherent spatiotemporal modes from short, sliding windows of data, and patterns modes are revealed by unsupervised clustering. The gDMD clusters correspond to the average modes within a population and closely resemble canonical RSNs. The sDMD use only data from a single scan, calculating subject-level RSNs and their temporal dynamics simultaneously. When sDMD is used to extract subject-level Default Mode Networks from the data of a single scan, it does so more reliably than ICA, while also having unambiguous time resolution. Finally, we show that the extracted temporal dynamics are highly reproducible within subjects and between trials.

DMD extracts additional information beyond what we have fully considered within the scope of this work. By construction, each DMD has an intrinsic temporal frequency. We have made note of these frequencies (e.g. as labels within Fig. 2) but have drawn no conclusions from them beyond noting that they are plausible compared to the known literature (i.e. generally < 0.1Hz). However, the preliminary analyses in Appendix B suggest that networks may have distinct and reproducible frequency content which could be characterized by future studies. Similarly, it remains to be systematically investigated the phase information in each mode and how they relate to resting state network organization.

Beyond data that has already been pre-processed and mapped onto “grayordinate” cortical surface space, it may be desirable to analyze volumetric data directly. In principle, there is no reason why our DMD approach cannot be applied equally well to voxel data; nevertheless, the quality of the networks thus extracted would be unknown and must be thoroughly characterized. Similarly, care should be taken when applying our approach to data acquired at different spatial and temporal resolutions. We have made all of our code openly available in the hope that these analyses can be performed readily and reproducibly.

This demonstration that DMD is capable of extracting modes reliably from short windows of data, with similar or better performance than ICA, lays the groundwork for future methodological developments. Indeed, a key motivation for the use of DMD is that a number of powerful extensions to DMD have been developed that could be readily applied to BOLD fMRI data. DMD with Control (DMDc [61]) uses a control theoretic framework, allowing intrinsic dynamics to be estimated from those driven by external stimuli. DMDc is a natural framework to analyze task-based fMRI scans. Multi-resolution DMD (mrDMD [62]) improves the quality of the temporal information extracted over multiple timescales. These straightforward extensions of DMD to multi-resolution and control frameworks have no direct counterpart in ICA, giving DMD a critical advantage over ICA in future analyses of multi-scale signal extraction and control. Similarly, Optimized DMD [63] is an implementation of the DMD algorithm which is more computationally expensive, but significantly improves the precision of the extracted frequency information, which would enable the study of how characteristic frequencies may vary between networks, as suggested by the preliminary analyses of Appendix B.

It is increasingly clear that RSN dynamics are significantly altered by neurological disorders, and their impact on the spatial and temporal structures of RSNs may be diverse and highly individualized. Rather than searching only for patterns in RSNs strongly resembling those seen in the general population, it may be fruitful to explore methods that are agnostic to the average. We propose that DMD is an approach well suited for data-driven extraction of individualized structure and dynamics of networks from single scans, and we believe the development of the DMD family of methods opens new doors for exploring and characterizing whole-brain dynamics as captured by fMRI.

## 4. Methods

### 4.1. Human Connectome Project rs-fMRI Data

We used rs-fMRI data from 120 unrelated individuals in the Human Connectome Project (HCP) dataset as provided by the WU-Minn Consortium [54, 55]. The HCP acquired four separate resting state scans using a multiband pulse sequence (multiband factor = 8, FA = 52°), each scan having 1200 time points (14.4 minutes, TR: 720ms, TE: 33ms, FOV: 208mm x 180mm, Matrix: 104 x 90, with 72 slices) [56]. For each scan, an equal number of volumes with left-right and right-left phase encoding directions were acquired. To correct the fMRI scans for gradient distortions, the HCP also acquired two spin echo EPI images with reversed phase encoding directions, with a unique pair of spin echo EPI images for each of the resting state acquisitions [64]. These spin echo EPI images were then used to accurately spatially normalize the fMRI volumes to the structural scans. For the remainder of this document, we label and refer to the four resting-state runs as “REST1_LR”, “REST1_RL”, “REST2_LR”, and “REST2_RL”).

The resting state scans had already been preprocessed by the HCP Consortium using the HCP minimal preprocessing pipeline [56]. In addition to gradient distortion correction, this preprocessing includes fMRI denoising using FIX [65], masking, motion correction, registration and interpolation of the timeseries onto the cortical surface (the CIFTI “grayordinate” space). Included in this preprocessing was the application of FreeSurfer to generate mesh representations of the the cortical surface. These FreeSurfer surface meshes were upsampled to ~164k vertices, and subsequently downsampled to ~32k vertices, resulting in an average vertex spacing of roughly 2mm at the cortical midthickness level. The resting state time series were then mapped onto this low-resolution 32k vertex mesh. This surface-mapped data consists of 91,282 “grayordinates” (~60k surface points mapping the cortical surfaces, along with ~30k subcortical voxels). In this manuscript, we restrict our attention to the 2*D* cortical surface mesh. Note, however, that DMD could be similarly applied to 3*D* voxelized data (see Discussion).

All data used in this manuscript are freely available for download from the HCP. Additionally, scripts for downloading the correct data, running all analyses described within this paper, and figure generation are openly available at https://github.com/kunert/DMD_RSN.

### 4.2. Dynamic Mode Decomposition

This section describes the specific DMD algorithm implemented in this manuscript [49, 50]. Each scan is broken into windows of data, and data from each window is collected into a data matrix **X,** where each row represents one of the *n* grayordinates and each column 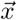 is one of the *m* time snapshots in that particular window:
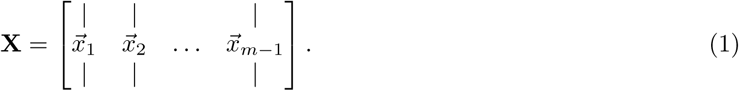

Here 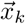 is a vector in grayordinates space giving the BOLD signal at time index *k.* Next, we also define the time-shifted data matrix **X′,** which is defined similarly but with each column advanced forward by a single timepoint:
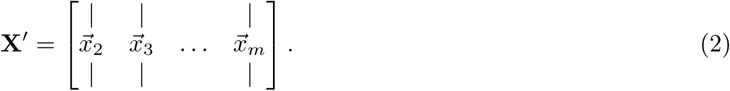

The goal of DMD is to describe the matrix **A** that best maps 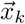 onto 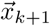, namely by solving for the eigenvalues and eigenvectors of **A.** In other words, we treat dynamics of the system as approximately linear and seek the **A** that best solves
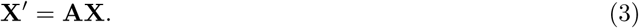

However, this **A** matrix is a *n* × *n* square matrix; for the HCP data, *n* = 91,282 grayordinants, so **A** has approximately 10^10^ elements — this is a tremendously large number of unknowns, and they are poorly constrained by the limited data in **X** and **X**′. Further, we hypothesize that many of these brain areas have strong correlations, so their dynamics are relatively low rank and may be explained by far fewer modes.

The DMD algorithm takes advantage of the singular value spectrum (SVD) of the data matrix **X** to obtain the dominant eigendecomposition of **A** without actually computing **A.** We first take the singular value decomposition (SVD) of the data matrix **X** = **UΣV*,** which decomposes the **A** into the product of unitary matrices **U, V** and diagonal matrix **Σ.** Because the singular values and singular vectors are ordered by decreasing energy, this decomposition can be used truncated by taking the first *r* columns of **U** to form a *n* × *r* matrix **U_r_.** We may similarly form a *r* × *r* matrix **Σ_r_** and a *m* × *r* matrix **V_r_.**

This truncation gives the optimal rank-r reconstruction of the data matrix, 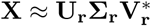. The choice of the number of SVD modes *r* is equal to the resultant number of DMD modes *r* and is used as a parameter throughout this manuscript.

Taking the SVD allows us to then calculate the pseudoinverse 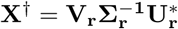 and simply solve for **A:**
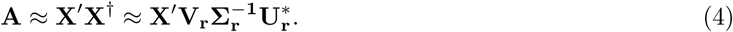

Given that the number of grayordinates is large, we do not consider the *n* × *n* **A** matrix directly, but instead project into the reduced-dimensional **U_r_** basis:
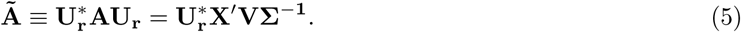

Additionally, we scale each mode according to how strongly it is present in the original data, and each modes is scaled by the singular values **Σ_r_** as in [50]:
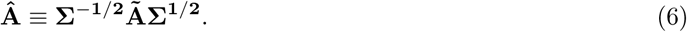

We then compute the eigendecomposition of this scaled, reduced dimensional 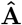 matrix:
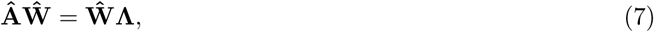

where **Λ** is the diagonal matrix of eigenvalues *λ_j_,* and the columns of 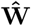 are the eigenvectors of 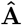. These eigenvectors can be used to compute the dynamic modes:
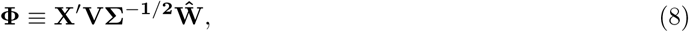

where the *j*-th column of **Φ** is the dynamic mode 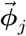. From the dynamic modes and corresponding eigenvalues, we can approximate the dynamics of the system as:
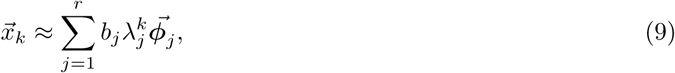

where the weights *b_j_* can be fit as initial conditions for at *k* = 0. We can also write this in terms of a continuous time *t* by writing the dynamics in terms of a complex exponential:
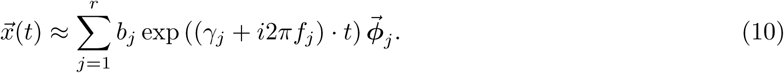

In the exponent, *γ_j_* is the real-valued growth/decay constant, and *f_j_* is the real-valued oscillation frequency of the mode in cycles per second (hertz). These are calculated from the real and imaginary components of the corresponding DMD eigenvalues:
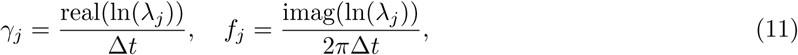

where Δ*t* is time between measurements, in seconds. This change of units does not carry any additional information over the complex eigenvalue *λ_j_,* but *γ_j_* and *f_j_* are readily interpretable as standard growth/decay constants and oscillation frequencies.

Like the eigenvalues, the dynamic modes 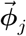 can be complex valued. In other words, each element of 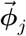 has both a magnitude and a phase. In this work, we consider only the magnitude of the dynamic mode 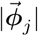; generally, when we refer to the “dynamic mode”, or when we do calculations such as spatial correlations, etc., we are referring to and using |*ϕ_j_*|. However, the complex-valued *ϕ_j_* also contains the relative phase between regions, which is of potential interest for future analyses. For real-valued data 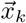, oscillatory modes appear as conjugate pairs 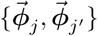 with the same spatial magnitudes |*ϕ_j_*| = |*ϕ_j′_* |; thus the number of unique spatial patterns extracted from a data matrix will be ≤ *r.*

### 4.3. Independent Component Analysis

fMRI data is often processed using Independent Component Analysis (ICA) [66]. Just as above, if we collect all data from a scan into a data matrix **X,** then we can formulate ICA as an attempt to solve the following:
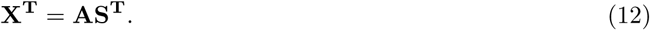

The goal of spatial ICA is to decompose the data into non-orthogonal, statistically independent source signals, which are the columns of the *n* × *r* matrix **S.** The *m* × *r* mixing matrix **A** gives the mixture of these signals, at each timepoint, which best approximates the original data.

However, because individuals subjects have their own unique time courses, comparing ICA components estimated at the subject-level is difficult, and consequentially, how to make inference about group-level components is not immediately clear. As an alternative, fMRI analyses often employ multi-subject ICA methods to estimate 1) population-based components, or 2) single-subject components informed at the group level. A variety of multi-subject methods have been developed, including temporal or spatial concatenation group ICA approaches [67, 68], higher-order tensor decomposition methods [40], and dual regression-based approaches [69, 70].

In this analysis, we use the FastICA algorithm as implemented in scikit-learn 0.18 [71]. FastICA attempts to maximize the statistical independence of the signals by maximizing the non-Gaussianity of the projected data, as detailed in [72]. In order to directly compare DMD to ICA, we compare both algorithms at two spatial component levels: using components informed at the single-subject level and using components informed at the group level. For the single-subject-level comparison, we compute single-subject ICA components using sklearn’s FastICA. For the group-level comparison, we use the group ICA and dual regression results generated previously by the HCP using Melodic’s Incremental Group-PCA algorithm (MIGP) [73]. MIGP is an iterative approach that incrementally incorporates time-series data for single subjects and updates a current running estimate of the set of spatial eigenvectors that best describes the time series data for the current set of incorporated subjects. MIGP generates a very close approximation to group-ICA components generated by classic temporal concatenation group-ICA approaches, but without the large computer memory requirements. The HCP then used dual regression to regress each individual subject’s time-courses on the MIGP-generated group-ICA components.

### 4.4. Sliding Window Mode Calculations

The rs-fMRI scans analyzed each consist of 1200 individual timesteps, which we break up into sliding windows. We use a simple square window (as opposed to a scheme which weights timepoints depending upon their position in the window), such that our calculations are performed on a submatrix of the full data matrix.

There are three parameters to consider when computing DMD and/or ICA modes within a sliding window: *T,* the length of each window; *dT,* the number of timesteps by which to slide the window; and *r,* the number of modes to compute within each window. We observed that our results are relatively robust to these parameters. Within the results shown in the main manuscript, we have chosen *T* = 32, *dT* = 4, and *r* = 8. In seconds, this corresponds to roughly 22.4s windows which are translated in increments of about 2.8s. As discussed in Section 4.2, oscillatory DMD modes appear in conjugate pairs, and thus ≤ *r* = 8 modes will be extracted within each window. For example, the gDMD set used in Fig. 2 (the modes calculated from the REST1_LR scans of 120 subjects) consists of 160,756 total modes.

### 4.5. Mode Visualization

DMD modes are computed on the full 59,412 -dimensional grayordinate space. However, this space is unnecessarily large for visualization purposes, for which we primarily want to visualize the macroscopic structure of individual modes. It is also disadvantageous for the clustering of modes; we want to cluster modes based upon their overall structural similarity, and applying some method of spatial smoothing is helpful in minimizing spurious, noise-driven correlations. For visualization and clustering purposes, we therefore bin the modes, computing the average magnitude of grayordinates within a particular patch of space.

Each grayordinate can be mapped onto a *3D* coordinate in space. From this information, we first classify all grayordinates as on the left/right cortex and on the lateral/medial side. Each of these groups of grayordinates are then projected onto the sagittal plane and divided into 40 bins in each direction. This choice of bin granularity is arbitrary but was chosen heuristically to yield good visualization and clustering performance. Bins which contain no grayordinates are discarded, resulting in a total of 3856 bins.

Given a 59,412 -dimensional DMD mode *ϕ_j_*, we may average over the grayordinates within each bin to yield a 3,856-dimensional binned mode *m_j_*. All modes visualized within this paper have been binned in this fashion. Furthermore, modes have been binned before being clustered or correlated against each other, which was seen to substantially improve the performance of clustering.

### 4.6. Hierarchical Clustering

Having performed sliding window DMD mode extraction on the full set of 120 individuals, we wish to cluster modes based upon their spatial similarity on both the single-subject and whole-group level. In both cases we use the same hierarchical clustering pipeline, described as follows, applied to either the set of modes extracted from a single scan (the sDMD case) or from the full set of scans (the gDMD case).

A few pre-processing steps were found to be helpful in generating clean, robust clusters. First, modes are spatially binned as described in Section 4.5. This reduces the dimensionality of modes for clustering purposes, averaging over noise while preserving larger-scale spatial structure. We then wish to filter out modes which lack large-scale continuous spatial structure. A quick heuristic for the spatial continuity of a mode is calculated and thresholded upon as follows: for each binned mode, we count the number of bins with a z-score ≥ 2 which also have a neighbor with z-score ≥ 2. Thresholding on *c* keeps only those modes which possess a certain level of spatial continuity (e.g. in Fig. 2 we cluster only modes with *c* ≥ 25).

To group this filtered set of modes into a small set of interpretable clusters, we use hierarchical clustering as implemented in SciPy 1.0.0[58]. There are many metrics of spatial similarity which can be used in such a procedure. Qualitatively, we found the best performance (as indicated by the formation of tight, discrete clusters which group modes with similar large-scale spatial structure) by thresholding modes at a defined z-score, *z_t_,* to generate spatial masks (in Fig. 2 we use *z_t_* = 2.5). These masks were then clustered based upon the average spatial correlation within a group of modes.

We then form flat clusters that have a cophenetic distance of no greater than a defined threshold *t* (i.e. using the ‘distance’ threshold option of the `scipy.cluster.hierarchy` function `fclust`). Fig. 2 shows the clusters which are formed using *t* = 0.955, whereas Fig. 3 shows finer-grained subclusters formed using *t* = 0.950. The choice of *t* will affect the size and granularity of the cluster assignment, and can be varied to obtain smaller subclusters of any particular cluster. Note that in Figs. 2 and 3 we have only visualized clusters containing a minimum number of modes (≥ 400 in the gDMD case of Fig. 2, and ≥ 5 for the sDMD case of Fig. 3).

We compare these modes against a set of canonical RSNs, using the parcellation of Gordon et al. [57]. In Fig. 2(B) we show the correlation of each mode against each of these canonical masks, where the ordering of the modes has been defined by the hierarchical clustering. The clustering succeeds at grouping modes which resemble similar RSNs, and many of the clusters appear to correspond to distinct networks: for instance, the bottom-rightmost cluster (cluster number thirteen) strongly corresponds to the “SMmouth” network. When the average of the modes within each cluster is plotted in Fig. 2(D), it indeed has the expected spatial structure.

### 4.7. Subject-Level DMNs

The subject-level Default Mode Networks (DMNs) shown in Fig. 4 are found by calculating the sDMD clusters for modes from a particular subject, and taking the average of the cluster which most resembles the DMN. The optimal choice of *z_t_* and *t* will vary from scan to scan; we therefore perform the clustering procedure several times for values of *z_t_* ∊ (1.5, 3.0) and *t* ∊ (0.65, 0.98), saving the output cluster average which has the highest correlation with the gICA result. Though the choices of *z_t_* and *t* afford relatively little control over the final spatial structure of the result, we must be mindful of the fact that we are selecting the best-performing result out of several different runs. For a fair comparison, we afford ICA the same opportunity: allowing the number of modes to vary in the range *r* ∊ (8,10,12,16), calculating ICA multiple times, and saving only the result which correlates most highly with the gICA result. This was seen to be necessary, as the output of the ICA algorithm is not deterministic and will sometimes yield a low-quality result (often yielding correlations of ~ 0.1, as in the long tail of Fig. 4(B)).

Each of these approaches were compared against the group-ICA and dual regression approach, the results of which are calculated and provided for download by the HCP [70]. This approach first calculates groupICA, which is ICA performed on the temporally-concatenated full dataset of scans from 1200 individuals. This calculates high-quality, population-level averaged gICA modes. The gICA mode corresponding to the DMN is then regressed against the scans for a single individual, to calculate an approximate timecourse of DMN activation within a particular individual’s scans. There is then a second regression step of the timecourses of all spatial coordinates against the averaged-DMN timecourse. This yields a spatial map of coordinates which have similar time dynamics to the population-level DMN mode, which in effect yields a subject-level spatial map of the DMN. This results in clean, high-quality maps as seen in Fig. 4, but has the drawback of requiring a reference mask calculated from the entire set of population data.

Fig. 4(B) plots how strongly each subject-level ICA/sDMD DMN correlates with both (i) the full set of gICA modes, and (ii) the gICA mode calculated for the same subject. These are calculated by thresholding the modes at a z-value of *z* = 2 and then calculating the correlation coefficients between the thresholded masks.

### 4.8. Characterizing Cluster Dynamics

Assigning time dynamics to individual clusters (as in Fig. 3(B)) follows trivially from the windowing process: each cluster consists of a collection of modes, each calculated in a particular window corresponding to a particular time. The time dynamics of a cluster are simply given by the times of the windows in which its constituent modes were found. Notably, this gives a binary measure of mode activation (a cluster either contains a mode within a particular time window or it doesn’t). This is distinct from other methods of assessing the temporal dynamics of modes, such as the common technique of correlating a spatial mode with each frame of a scan; such techniques yield a continuous measure of mode activation, which does not fully disambiguate the activity of a mode from that of overlapping spatial patterns or random fluctuations. This disambiguation is accomplished in our approach through the assignment of modes into discrete clusters.

The gDMD pipeline was performed independently on the four sets of scans for all 120 individuals (the sets “REST1_LR”, “REST1_RL”, “REST2_LR”, and “REST2_RL”). This resulted in four sets of gDMD modes, which were correlated against each other to find the modes which were most similar across all sets. We chose the top six most consistent modes, as described in Appendix A, with the resulting modes visualized in Fig. 5(A). Each set of modes has associated time dynamics; as examples of how to characterize these dynamics we calculate both the occupancy matrix *O_ij_* and transfer matrix *T_ij_.* The occupancy matrix *O_ij_* indicates the number of windows in which both modes *i* and *j* are active. The diagonal elements *O_ii_* count the total number of times mode *i* was active (with or without the presence of other modes). The transfer matrix *T_ij_* shows the probability that if mode *i* is present, then mode *j* is present 30s later.

We assess the reproducibility of our approach by comparing the occupancies of each mode by each individual between different scans. This rests upon the assumption that if an individual shows increased activity in a particular mode in one scan, they are then more likely to be more active in the same mode within another scan. Note that we do not expect a perfect correlation near 1.0, but we do anticipate a significant one; for example, if an individual has a particularly active DMN in one scan, we may expect that that same individual is more likely to have an active DMN in another scan, but not that the DMN should be active for the exact same amount of time. An example of such a comparison is plotted in Figure 6, where each point represents a different individual, and we do indeed observe a positive correlation. We correlated all combinations of the four scans against each other for all different modes, and report the median correlation and occupancy within the table of Fig. 6. A significant correlation (in the sense that *p <* 0.05) is clear for all modes except for mode 4, indicating that our characterization of the dynamics indeed encodes underlying dynamic properties specific to different individuals.

## 5. Acknowledgements

The authors thank Nikita Sakhanenko for useful discussions. JMK and DJG acknowledge support from the Bill and Melinda Gates Foundation, and the Pacific Northwest Research Institute. JNK acknowledges support from the Air Force Office of Scientific Research (FA9550-17-1-0329). BWB acknowledges support from the National Science Foundation (award 1514556), the Alfred P. Sloan Foundation, and the Washington Research Foundation. SDR acknowledges support from the National Institutes of Health/National Institute on Aging (K01AG055669).

## Appendix A Comparing Clusters Across Scans

**Figure 1:**
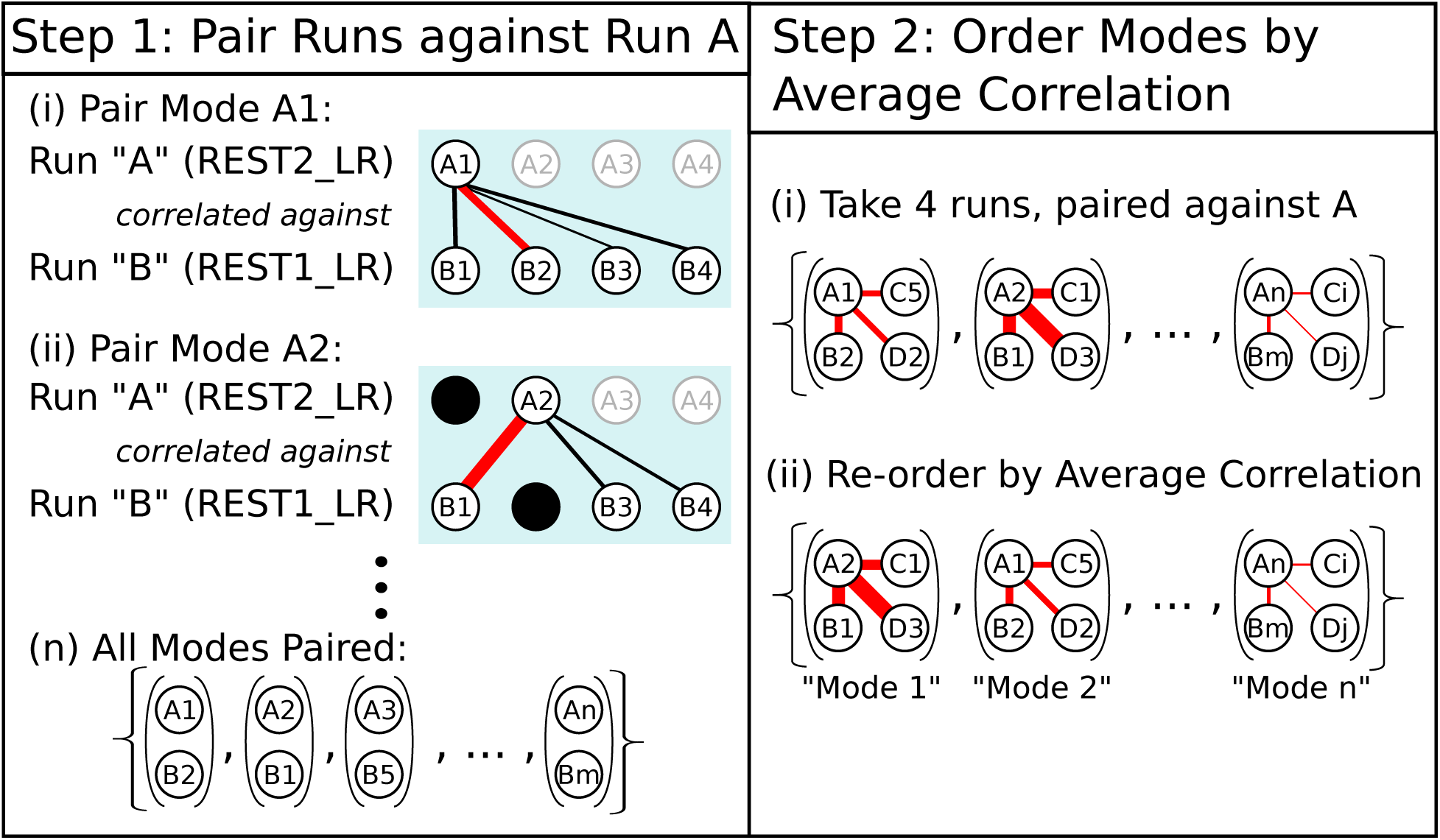
Procedure for pairing in ordering modes between different runs. When we process the four runs independently with identical parameters, they result in a different number of clusters, and thus a different number of corresponding averaged modes. Here we label the runs A, B, C, D; in the data, these refer to REST2_LR, REST1_RL, REST2_RL and REST1_LR, which yielded 36, 48, 49 and 42 modes respectively. We choose the run with the least number of modes (REST2_LR) as Run A, and match its modes to a unique mode from each of the other runs. **Step 1:** Iterate through the modes of Run A and match them (without replacement) to the most highly spatially correlated mode from Run B, until all modes are paired. Repeat this process for Runs C and D. **Step 2:** Calculate the average correlation between the A modes and the paired B/C/D modes, and re-order the modes by their average correlation. The modes which have the highest average correlation (i.e. which appear, on average, most similar between runs) we dub the “most consistent”. Spatial correlations of the B/C/D modes against the paired A modes in our results are plotted in Figure 2.

**Figure 2:**
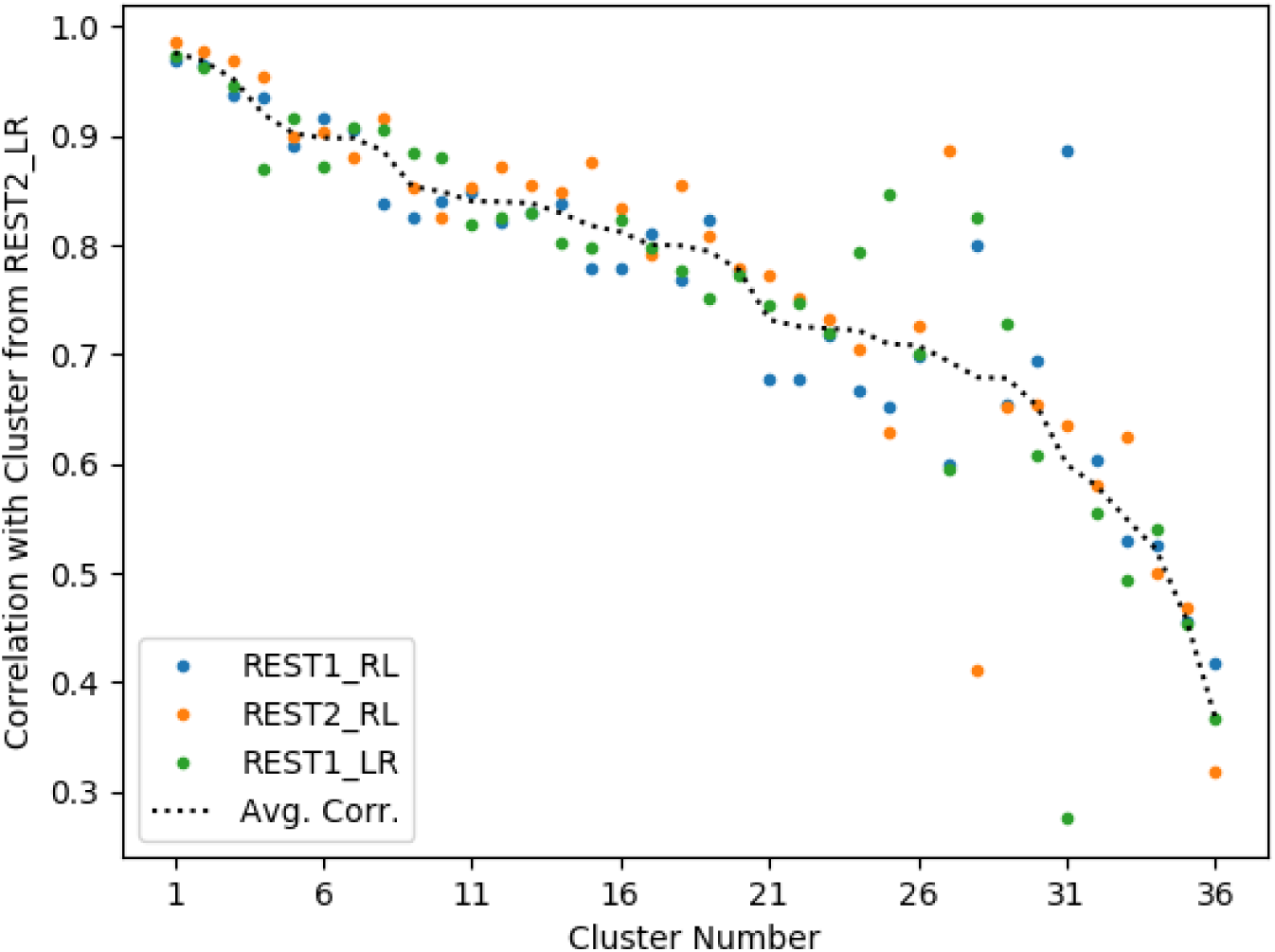
The averaged modes from each of the clusters from all four runs were paired together as described in Figure 1. As described there, these cluster-average modes are ordered by the average spatial correlation with those from the run REST2_LR. In the main manuscript, we visualize and analyze the dynamics of the top 6 modes, though this cut-off is arbitrary: similar spatial consistency appears in higher-numbered modes. For completeness, we include visualizations of the averaged modes of all clusters for all four runs on the following pages.

### Appendix A.1 Averaged Modes from All Clusters

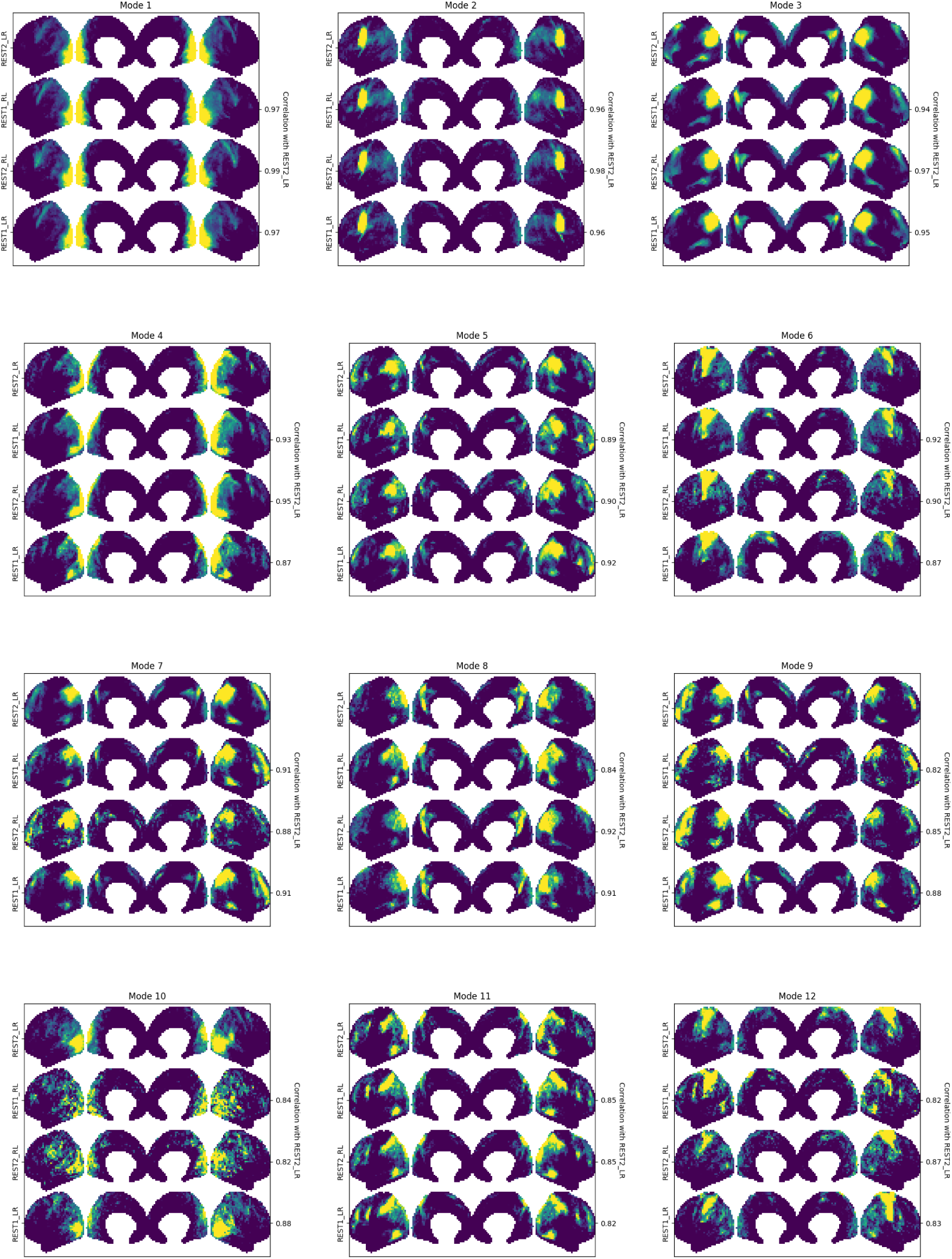
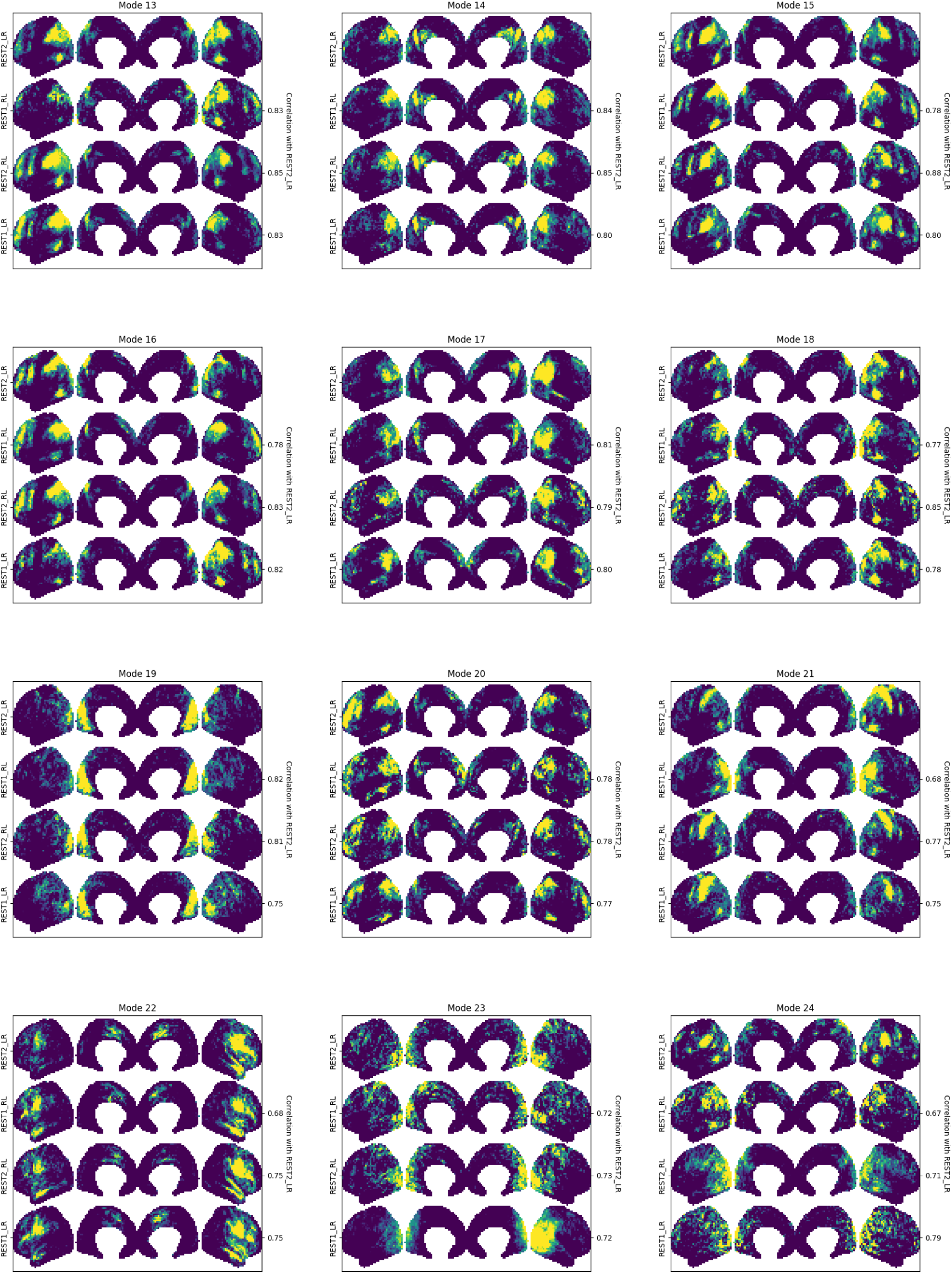
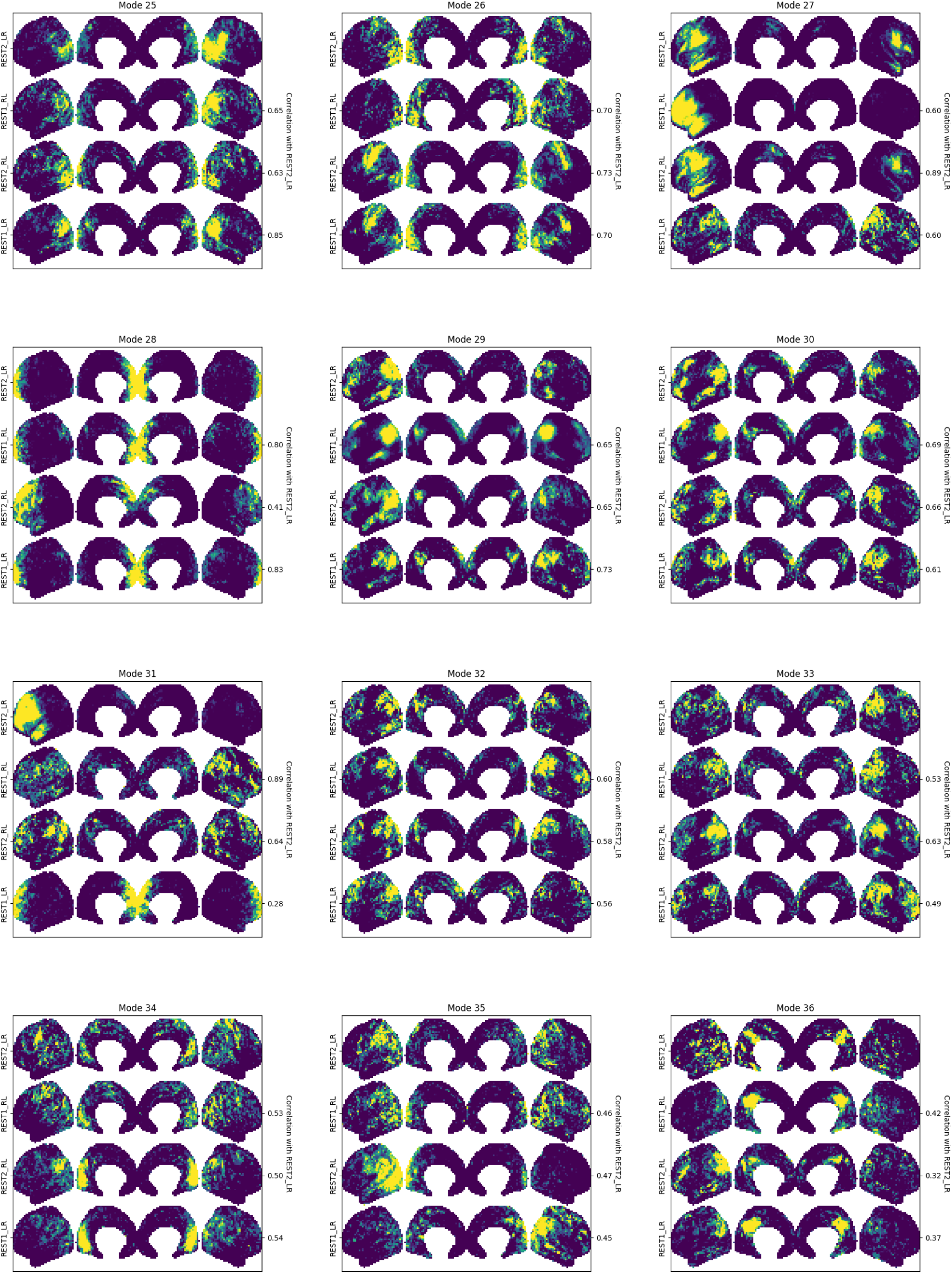

## Appendix B Cluster Frequency Content

In the previous section, we paired clusters extracted from four different runs and then averaged the spatial modes in each of the clusters. Spatially, the average mode of each cluster appears quite similar across the four different runs. DMD extracts a temporal frequency to accompany each spatial mode, and thus each cluster also has a distribution of frequencies. The distribution of frequencies in each cluster in each of the four runs in shown in Figure 1.

**Figure 1:**
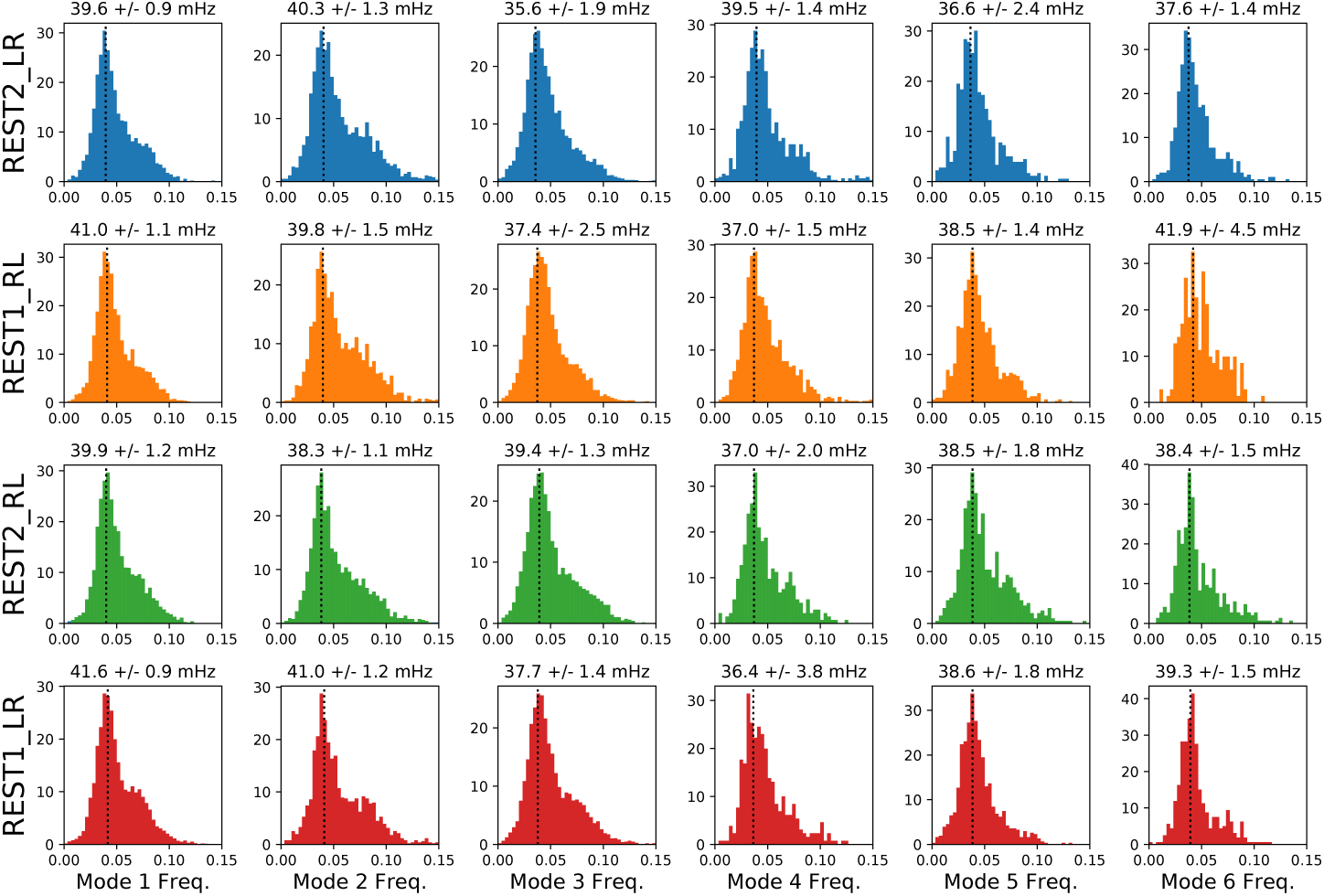
Frequency distributions for the modes constituting each of the six clusters in each of the four runs. Distributions appear approximately bimodal with a maximum-likelihood (peak) frequency at approximately 0.04Hz. The peak frequency (and its uncertainty) are approximated as described in this section, and are shown for each cluster.

The frequency distributions in Figure 1 are approximately bimodal; they are clearly not Gaussian, but appear to be well-approximated by the sum of two Gaussians, the first with a central frequency of ~0.04Hz and the second, shorter Gaussian with a central frequency of ~0.075Hz. It would be un-informative for these distributions to simply compute and compare the means. Instead, we fit a Gaussian Mixture Model to the frequency data using the `scikit-learn` module `mixture`. We select the optimal number of components using the Akaike information criterion, and take the maximum likelihood of the fit distribution. We perform this fitting procedure 100 times for each cluster on randomly-selected subsets of 75% of the data, and take the mean and standard deviation of these 100 trials as our estimate of the peak frequency and its uncertainty. This estimate of the peak frequency and its uncertainty are plotted for each mode in Figure 2

**Figure 2:**
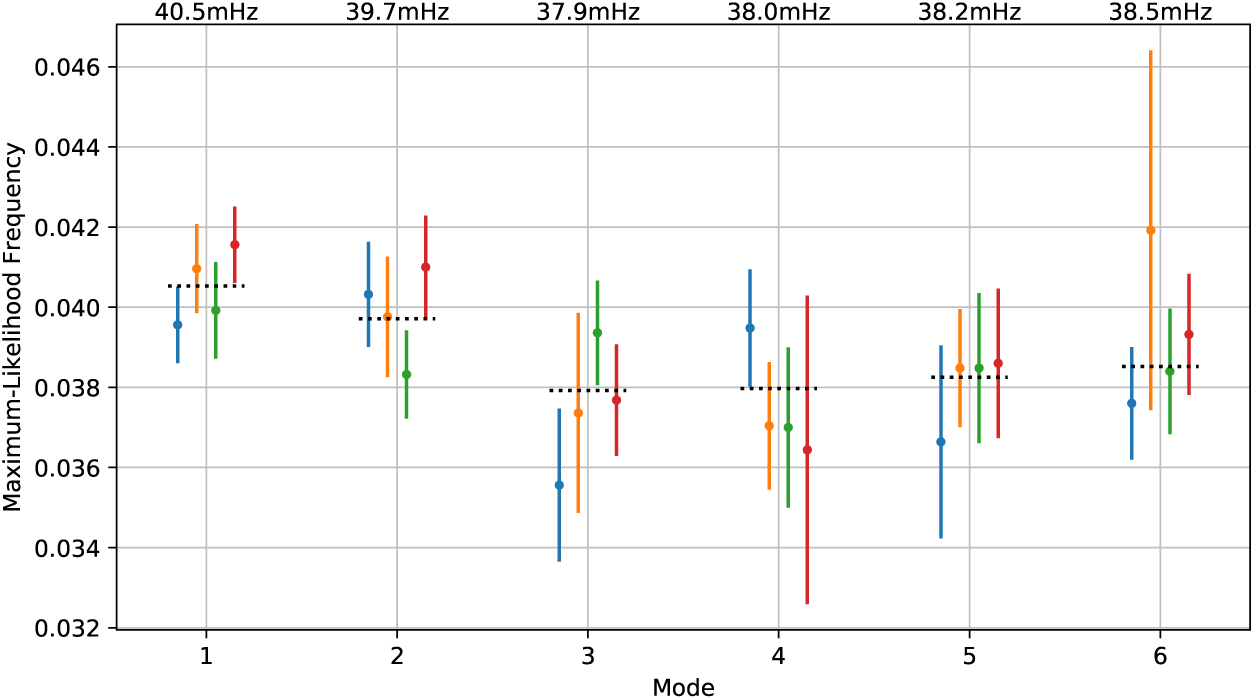
Peak frequencies inferred from the frequency distributions in Figure 1. For each mode, the peak was estimated for 100 random subsets of 75% of the data, and the resulting mean and standard deviation are plotted here. The dashed black lines indicate the mean frequencies of each cluster across the four runs. In short, the estimated peak frequencies are too imprecise to yield definitive conclusions, but these preliminary results suggest that the frequency content of different modes may vary reproducibly.

The distributions in Figure 1 are broad, and the estimates of peak frequency in Figure 2 are subsequently imprecise. This makes it difficult to make definitive statements about how the frequency content of the clusters are reproducibly varying between different scans. However, this preliminary analysis suggests possible reproducible variation: for example, Modes 1 and 2 appear to reliably have a peak frequency of close to 0.040Hz, whereas Modes 3-5 have a slightly lower peak frequency of 0.038Hz.

These preliminary analyses leave several open questions: does each network (corresponding to each cluster) have a distinct and reproducible frequency distribution? Does the peak frequency of each network vary from individual to individual, and could this individual variation in RSN frequency serve as a useful biomarker? In this paper we use the “exact DMD” algorithm for computing DMD, which is computationally efficient but yields a biased and imprecise estimate of the frequency. This has led to methods such as Optimized DMD ^1^, which considerably improves the precision of the frequency estimate but is much more computationally expensive. Future work should consider the use of these more expensive methods, as they may be able to characterize RSN frequency to a level of precision such that RSN frequency variations could be characterized down to the level of single subjects.

## Appendix C Singular Value Spectrum

In calculating windowed DMD, we need to select the number of modes to extract from each window. Throughout the paper we have chosen *n* = 8 modes to be extracted from each of our 32-frame windows. The appropriateness of this choice can be investigated by looking at the distribution of singular values. In this section, we look at the singular values from all the windows from a single scan (Subject 102513, run REST1_LR). DMD is performed over 32-frame windows, slid over a 1200-timestep scan in increments of 4-frames, leading to 293 windows. DMD is thus performed 293 times, including 293 SVD computations. The distributions of singular values from each of these computations is shown in Figure C1.

**Figure C1:**
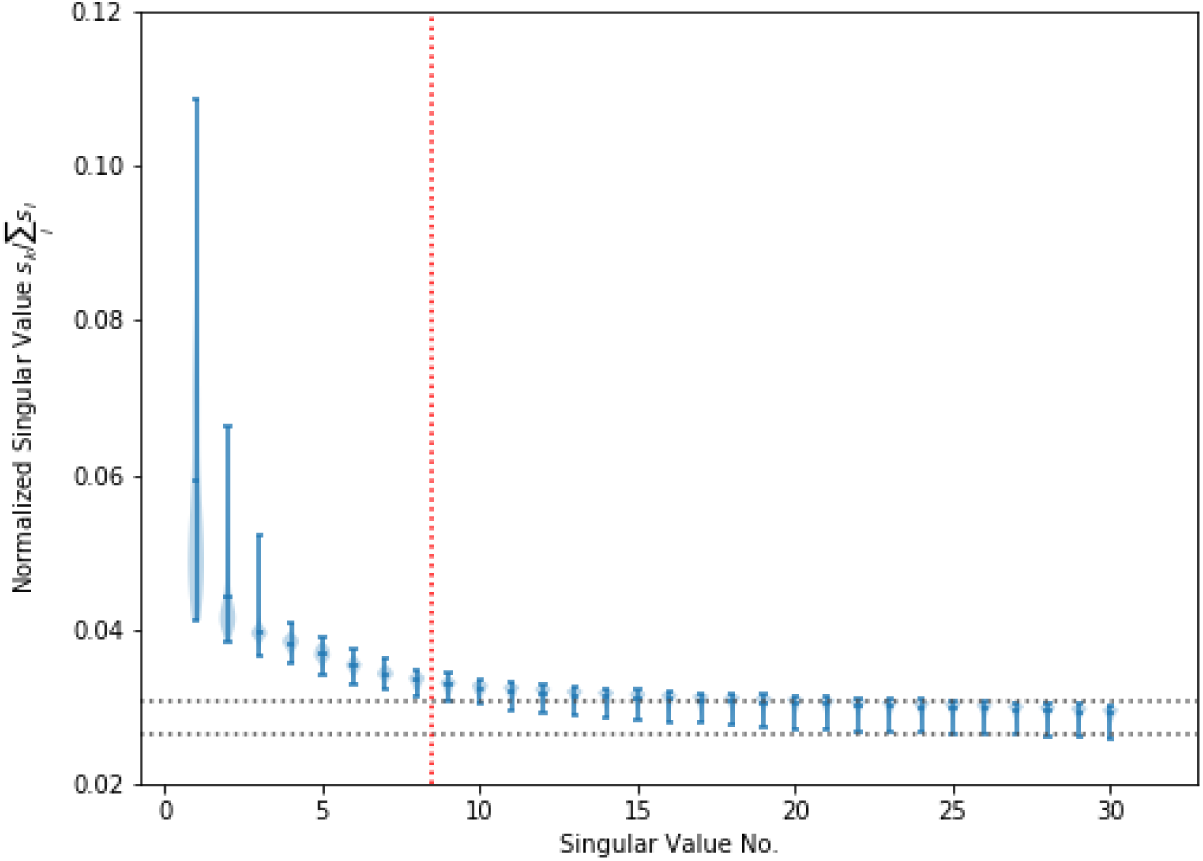
Distributions of singular values over the 293 windows of a single scan (Subject 102513, run REST1_LR). Our cutoff includes the first 8 modes (indicated by the dotted red line), which sums to only ~ 40% of the energy as there is a large amount of noise. We roughly approximate a “noise floor” for singular values by taking the average minimum/maximum singular values of Modes 20-30, yielding the range indicated by the black dotted lines in the figure. Figure C2 shows a zoomed version of this figure, to better display the distributions close to our *n* = 8 cutoff.

**Figure C2:**
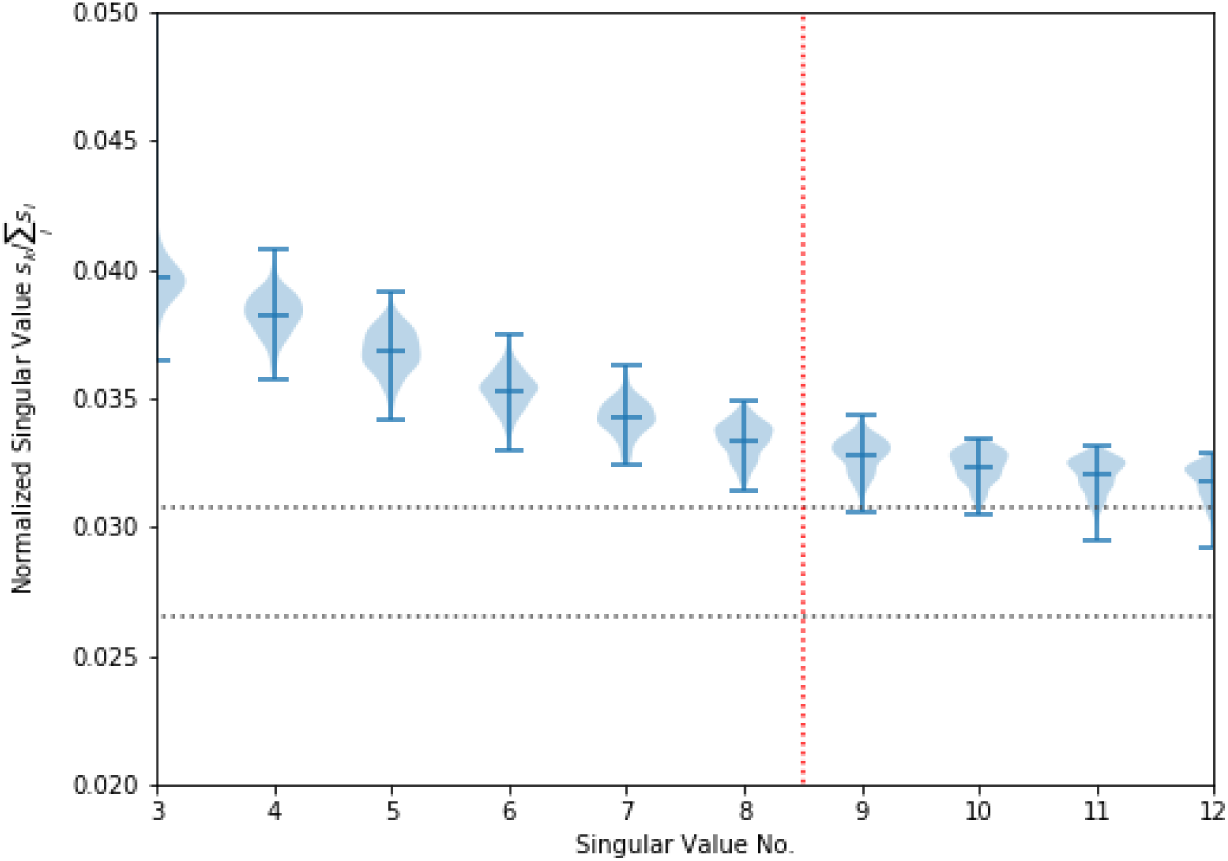
The same as Figure C1, zoomed in around the *n* = 8 cutoff. After this cutoff, singular values begin to dip into the “noise floor” range. This suggests that *n* = 8 is a reasonable choice, but also suggests that similar nearby choices of *n* should yield similar results, as modes in this range carry only a small (though potentially non-negligible) amount of energy. Indeed, we find that our results are not particularly sensitive to the precise choice of *n.*

## Footnotes

1 T. Askham and J.N. Kutz. “Variable projection methods for an optimized dynamic mode decomposition.” arXiv preprint arXiv:1704.02343,2017.

